## Supplementary figures for "Neurons in the bat auditory cortex encode class and complexity of future vocalizations"

### Supplementary materials

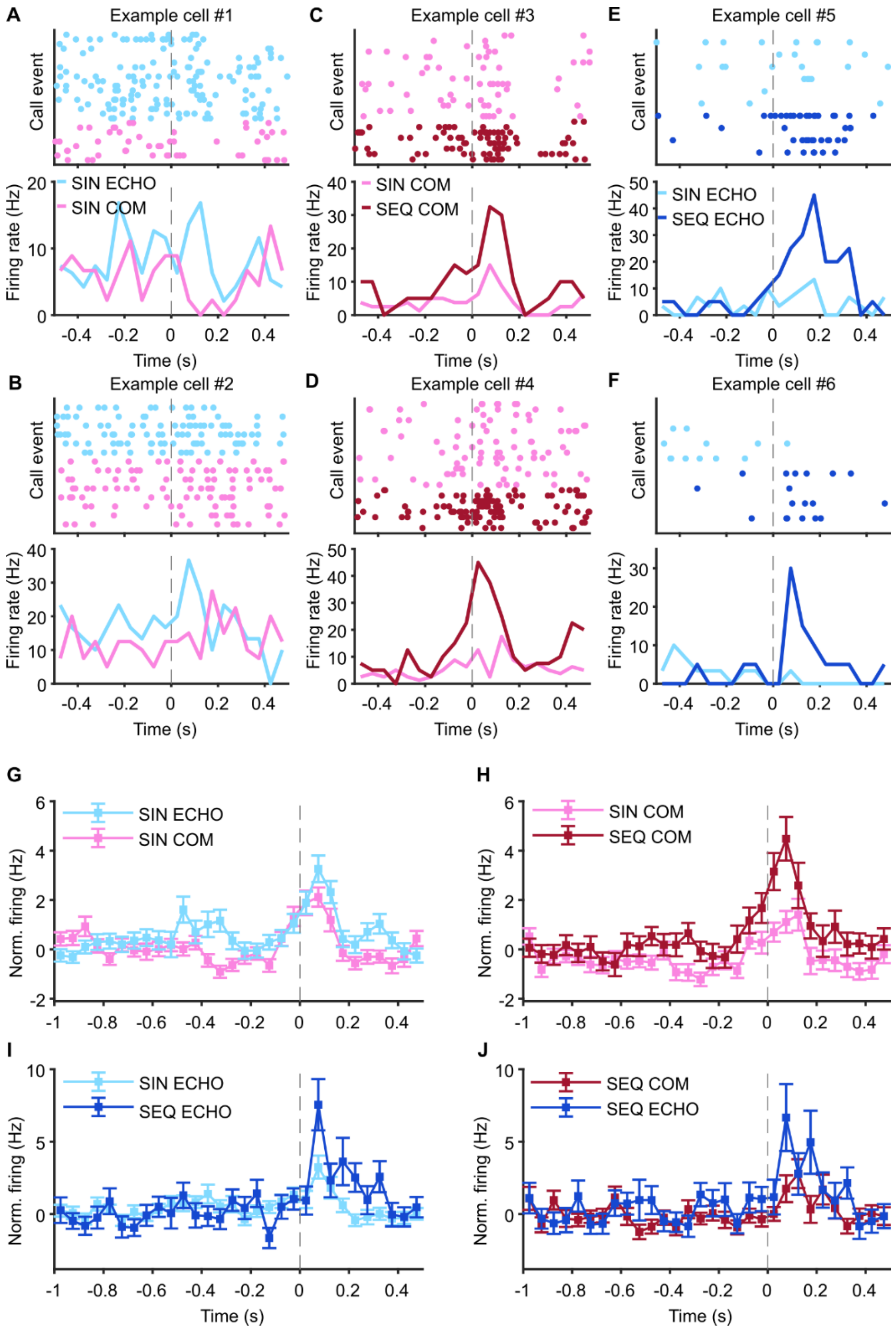


**Figure S 1:** Example neurons and extended pre-time for neuronal firing rates to different call categories, related to Figure 2. A-F, Example cells recorded during SIN ECHO and SIN COM (A-B), during SIN COM and SEQ COM (C-D), or during SIN ECHO and SEQ ECHO (E-F) with raster plot (top) and averaged firing rates (bottom). G-J, Averaged firing rates as in Figure 2 showing activity 1 s prior to vocal onset.

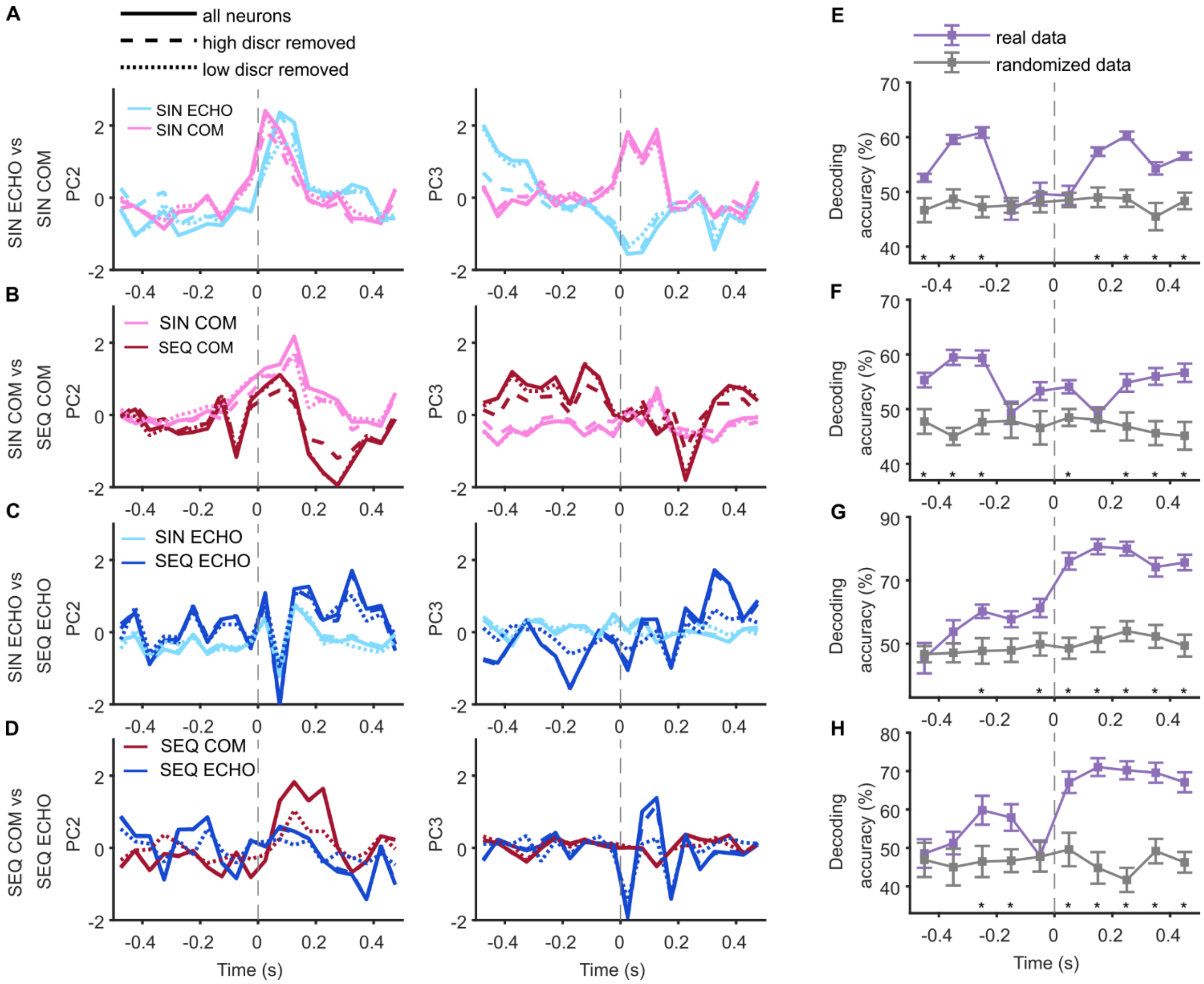


**Figure S 2:** Additional analysis using neuronal activity to different call categories, related to Figure 2. A-D, 2nd (left) and 3rd (right) PC scores over time from PC analysis on firing rates to call categories of neurons in Figure 2. Solid lines indicate dataset with all neurons, dashed lines indicate dataset without high discriminator neurons (30% of neurons with highest modulation indices), dotted lines indicate dataset without low discriminator neurons (30% of neurons with lowest modulation indices). E-H, SVM decoding of call category using neuronal activity vectors to all call events from all recorded neurons. *p<0.01 for accuracy using real firing rates vs. randomized dataset, Wilcoxon signed-rank test.

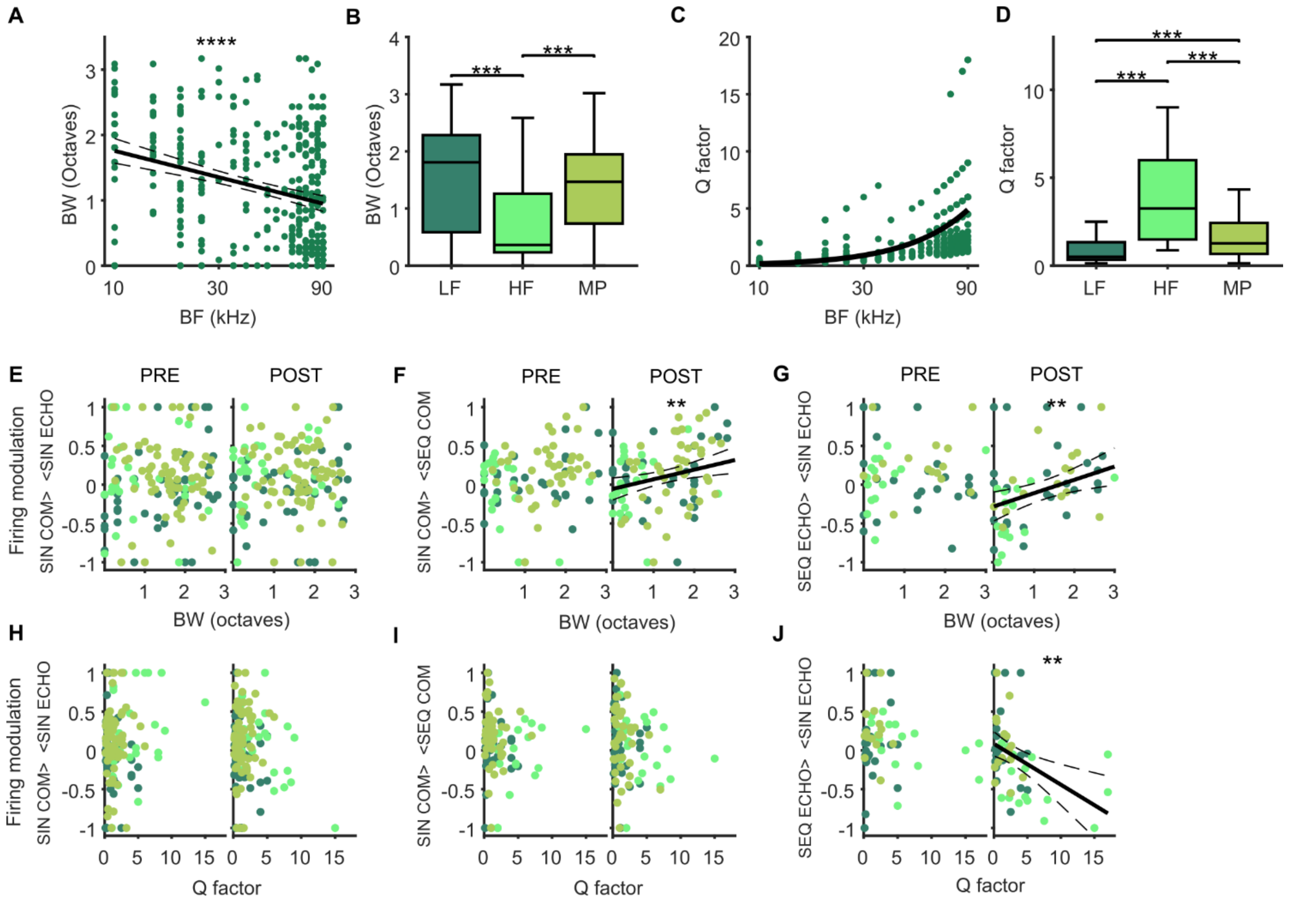


**Figure S 3:** Additional analysis of frequency tuning properties and relation to call modulation, related to Figure 4 and 5. A, BF and respective BW of every neuron. Solid line indicates linear regression fit, dashed lines indicate 95% CI, asterisks indicate significance of linear regression model with ****p<0.0001. B, BW in LF, HF and MP neurons. C, BF and Q-factor of every neuron. Solid line indicates exponential curve fit. D, Q-factor in LF, HF and MP neurons. Box plots in (B) and (D) represent median (line), 25$^{th}$ and 75$^{th}$ percentile (box) and whiskers extend to the minimum and maximum values within 1.5 times the interquartile range. ***p<0.001, Wilcoxon rank-sum test. E-J, Vocalization-dependent firing modulation index pre (left) and post (right) vocal onset and the respective BW (E-G) or Q-factor (H-J) of each neuron. Modulation indices for SIN ECHO vs. SIN COM (E, H), for SEQ COM vs. SIN COM (F, I) and for SEQ ECHO vs. SIN ECHO (G, J). Solid line indicates linear regression fit, dashed lines indicate 95% CI, asterisks indicate significance of linear regression model with **p<0.01.
